# Nuclear lamin A-associated proteins are required for centromere assembly

**DOI:** 10.1101/2023.09.25.559341

**Authors:** Adriana Landeros, Destiny A. Wallace, Amit Rahi, Christine B. Magdongon, Praveen Suraneni, Mohammed A. Amin, Manas Chakraborty, Stephen A. Adam, Daniel R. Foltz, Dileep Varma

**Affiliations:** Dept. of Cell and Developmental Biology, Northwestern University Feinberg School of Medicine, 303 E. Chicago Ave Chicago, IL 60611; Dept. of Biochemistry and Molecular Genetics, Northwestern University Feinberg School of Medicine, 303 E. Chicago Ave Chicago, IL 60611

**Author notes:** Contributed equally (Adriana Landeros, Destiny A. Wallace).

**Keywords:** Centromere, Mitosis, LaminA, Emerin, BANF1, CENP-A

## Abstract

Many Lamin A-associated proteins (LAAP’s) that are key constituents of the nuclear envelope (NE), assemble at the “core” domains of chromosomes during NE reformation and mitotic exit. However, the identity and function of the chromosomal core domains remain ill-defined. Here, we show that a distinct section of the core domain overlaps with the centromeres/kinetochores of chromosomes during mitotic telophase. The core domain can thus be demarcated into a kinetochore proximal core (KPC) on one side of the segregated chromosomes and the kinetochore distal core (KDC) on the opposite side, close to the central spindle. We next tested if centromere assembly is connected to NE re-formation. We find that centromere assembly is markedly perturbed after inhibiting the function of LMNA and the core-localized LAAPs, BANF1 and Emerin. We also find that the LAAPs exhibit multiple biochemical interactions with the centromere and inner kinetochore proteins. Consistent with this, normal mitotic progression and chromosome segregation was severely impeded after inhibiting LAAP function. Intriguingly, the inhibition of centromere function also interferes with the assembly of LAAP components at the core domain, suggesting a mutual dependence of LAAP and centromeres for their assembly at the core domains. Finally, we find that the localization of key proteins involved in the centromeric loading of CENP-A, including the Mis18 complex and HJURP were markedly affected in LAAP-inhibited cells. Our evidence points to a model where LAAP assembly at the core domain serves a key function in loading new copies of centromeric proteins during or immediately after mitotic exit.

## Results and Discussion

The kinetochores of the chromosomes must attach effectively to spindle microtubules to ensure proper separation of sister chromatids during mitosis. Kinetochores assemble at the centromeric regions of chromosomes during mitosis and serve as attachment sites for microtubules of the mitotic spindle. The protein CENP-A, a Histone H3 variant, replaces Histone H3 at the centromeric nucleosomes to define the centromeric region of the chromosomes. The deposition of CENP-A epigenetically controls the assembly of the kinetochores during mitosis [1, 2]. In addition, histone adaptors and chaperones, including the Mis18 complex and HJURP, are critical for new CENP-A deposition at pre-existing centromeres [3, 4]. Kinetochores are composed of the inner and outer kinetochore domains [5, 6]. The inner kinetochore domain consists of a conglomeration of protein complexes that organize into the constitutive centromeric associated network (CCAN). The CCAN is made up of 16 centromeric proteins (CENPs), including CENP-C and CENP-T, which function to connect the centromeric region to the outer domain of the kinetochore [1, 2]. The outer domain is composed primarily of the KMN protein network, which in turn consists of 3 protein complexes: Knl1, Mis12, and the Ndc80 complexes and constitute the core microtubule attachment site at kinetochores [5, 7].

### LMNA-associated proteins co-assemble with the centromeres at the outer core domain

Following anaphase chromosome segregation, as a part of the early events in nuclear envelope reformation, many components of the nuclear lamina undergo dephosphorylation and assemble at chromosomal regions referred to as the core or non-core domains in telophase [8]. It is known that LMNA/C and its associated proteins (LAAPs), including BANF1/BAF and Emerin (EMD), assemble laterally on each side of the segregated chromosomes at chromosomal regions referred to as the outer and inner core domains, as shown by the schematic representation (Figure 1A) and the associated images (Figure 1B) [9]. A previous study has shown a colocalization between BANF1 and the inner kinetochore protein, CENP-C, in *Drosophila* S2 cells that was important for centromere function [10]. To test whether this colocalization also occurred in human cells, we immunostained for centromere/kinetochore markers, including CENP-A, CENP-C and CENP-T in a HeLa stable cell line expressing GFP-BANF1. We did not observe any colocalization between centromeres/kinetochores and BANF1 or any other LAAPs during interphase (Figure S1A) or any early stages of mitosis including anaphase (not shown) in multiple human cell lines. However, interestingly, we found that LAAPs do colocalize to a substantial extent with centromeres at the region corresponding to the outer core domain proximal to the location of the mitotic spindle in telophase cells. For example, we found that centromeric marker, CENP-A and inner kinetochore protein CENP-C both colocalized with GFP-BANF1 at the outer core domain of the segregated chromosomes during telophase (Figure 1C). Further, we also found that CENP-A, CENP-C and CENP-T colocalize with EMD at the same domain during telophase in HeLa cells (Figure S1B). Additionally, we found an overlap between the inner kinetochore marker, CENP-T and EMD at the outer core also in RPE1 cells (Figure 1C). In this study, we refer to this part of the core as the kinetochore-proximal core (KPC) domain to demarcate it from the other part of the core that is adjacent to central spindle microtubules (kinetochore-distal core or KDC), which did not show overlap with the kinetochores/centromeres (Figure 1A).

**Figure 1.**
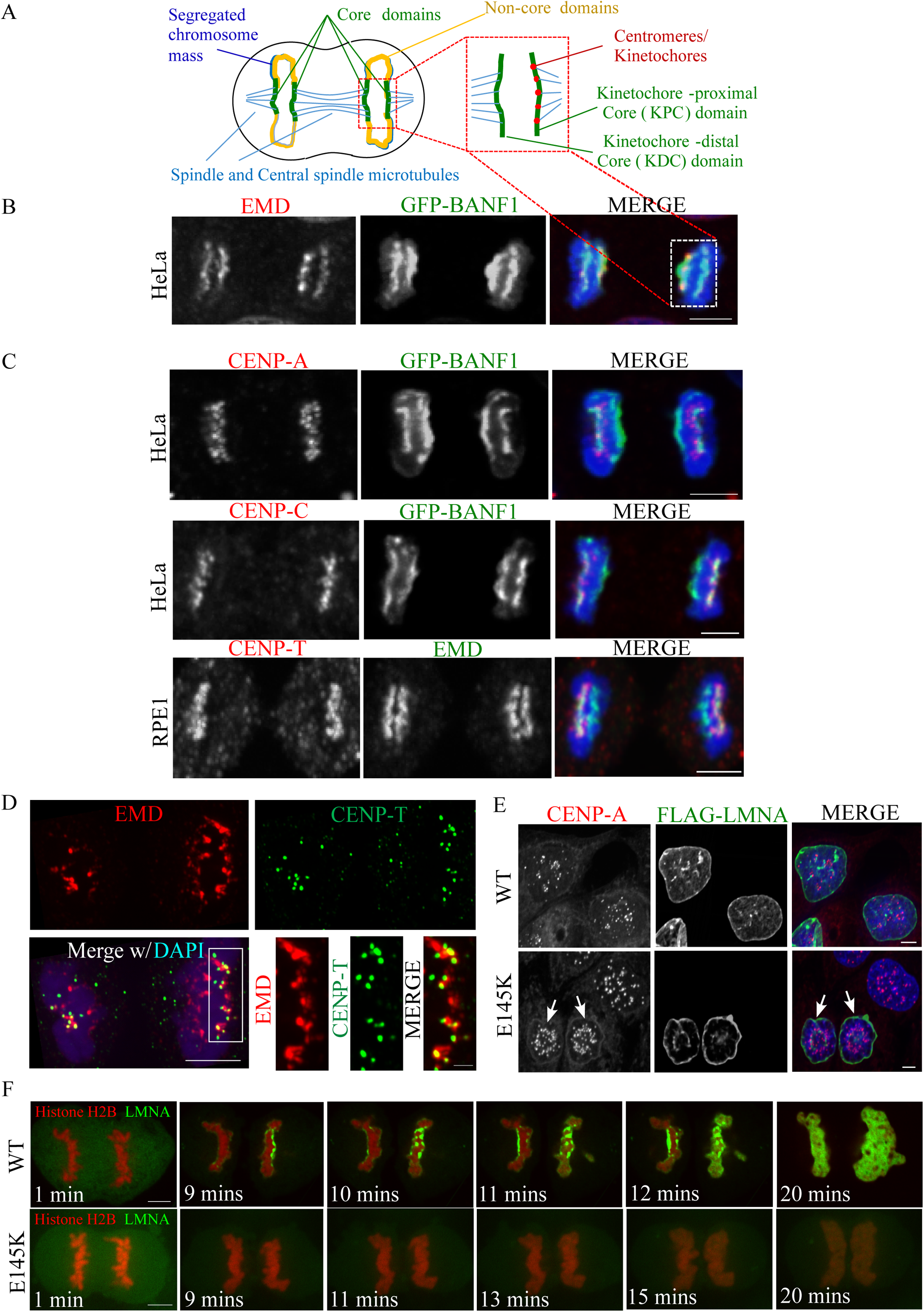
Lamin A-associated proteins (LAAPs) co-assemble with centromeric components at the kinetochore-proximal core (KPC) during telophase. (A) A schematic representation of the chromosomal core domains formed during mitotic telophase. The core domains assembled on the segregated chromosome mass are indicated in green, the non-core domains in yellow, the spindle and central spindle microtubules in blue and the centromeres/kinetochores in red. (B) Visualization of the core domain via the immunostaining of LAAPs, BANF1 (GFP, green) and EMD (red), with the chromosomes counterstaining using DAPI (blue). The selected regions from the schematic on top and the mitotic cell at the bottom indicate the two core domains formed in telophase. (C) Immunostaining of CENP-A combined with GFP-BANF1 in HeLa cells (top panel); CENP-C with GFP-BANF1 in HeLa cells (middle panel) and CENP-T with EMD in RPE1 cells (bottom panel). Scale bar, 5 µm. (D) HeLa cells in telophase immunostained using anti-EMD and anti-CENP-T antibodies with the chromosomes counterstained using DAPI, were subjected to Structured-Illumination (SIM) super-resolution imaging. Bar, 5 µm. Insets from C are shown at the bottom right. Bar, 1 µm. (E) HeLa cells expressing FLAG-tagged WT or E145K mutants of LMNA were immunostained for the FLAG-tag and the centromeric protein, CENP-A, with the nuclei counterstained using DAPI. (F) Live imaging of HeLa cells stably expressing Histone H2B-RFP transfected with either GFP-tagged WT (upper panel) or E145K LMNA (lower panel) constructs. Bar, 5 µm.

To derive a clear understanding of the spatial arrangement of centromeres and LAAPs at the KPC, we utilized the super-resolution imaging technique, Structural Illumination Microscopy (SIM). Using SIM acquired images of HeLa cells immunostained with antibodies against EMD and the CCAN (inner kinetochore) component, CENP-T during telophase, we found that CENP-T is embedded into or colocalized with larger plaque like structures formed by EMD (KPC), while certain regions of the KPC showed no overlap with the inner kinetochores (Figure 1D). Together these observations lead us to hypothesize that (i) LAAPs play a critical role in centromere assembly and/or that (ii) centromeres may be providing a framework for LAAPs to populate at the KPC domain.

### Perturbation of LMNA function interferes with centromere assembly and distribution

LMNA/C function is known to be important for nuclear architecture and chromosome organization. Mutations in the *LMNA* gene cause defects in the nuclear envelope, a common phenotype in children with Hutchinson-Gilford progeria syndrome (HGPS), an extremely rare, progressive genetic disorder that causes children to age rapidly [11]. Interestingly, previous studies have shown that in dermal fibroblasts from patients with a mutation (E145K) in the *LMNA* gene, in addition to the expected defects in the nuclear morphology, there was also an abnormal distribution/clustering of centromeres, marked by the anti-CREST antiserum (ACA), towards the center of the nucleus [12]. Similar results were observed by live imaging of HeLa cells stably expressing YFP tagged CENP-A, transfected with either FLAG-tagged WT or E145K LMNA constructs. We confirmed this previously observed phenotypes of abnormal nuclear morphology and centromeric clustering in HeLa cells by transfecting with a FLAG-tagged WT LMNA or the E145K mutant and carrying out immunostaining for the *bona fide* centromeric marker, CENP-A (Figure 1E). In addition to their altered distribution, the centromeric foci were also visibly smaller in LMNA E145K expressing cells (Figure 1E, arrows) [12], even though the published study did not perform a further detailed analysis of this phenotype.

It is known that LAAPs are deposited at the core domains in telophase, which is one of the early stages of the nuclear envelope reformation [8, 13–15]. For example, previous work has established the time at which BANF1 and EMD assemble at the core domain of chromosomes. Additionally, it is known that BANF1 localization to the reforming nuclear envelope in telophase is critical in recruiting EMD [14]. We validated this data for the timing of BANF1 at the core domains via live cell imaging of GFP-BANF1 HeLa cells (Figure S1C; Video S1). Further, live cell imaging of HeLa cells stably expressing Histone H2B-RFP transfected with either GFP-tagged WT (Video S2) or E145K LMNA (Video S3), demonstrate that WT LMNA localizes to the core domain immediately after BANF1 and EMD localization while the mutant LMNA does not deposit properly at the core domains (Figure 1F; Video S2 & S3). From the observations so far, we hypothesize that the co-assembly of the LAAPs and the centromeres at the core is important to maintain the structural identity of the KPC, which in turn has a critical role in centromere assembly and function.

As mentioned, variations in centromere assembly and distribution was reported in human embryonic fibroblasts or HeLa cell nuclei expressing the E145K mutation in the *LMNA* gene [12]. To analyze this phenotype in detail, we first treated HeLa cells with siRNA targeting LMNA (siLMNA) and measured the fluorescence intensity of inner kinetochore CCAN components, CENP-C and CENP-T, at kinetochores in both interphase and mitotic cells. We found that CENP-C and CENP-T kinetochore fluorescence is significantly reduced in both interphase as well as mitotic cells after siLMNA (Figure S2A-F). Next, we utilized a rescue approach to determine the effect of the E145K LMNA mutation on centromere and CCAN components. We generated a *LMNA KO* HeLa cell-line using CRISPR technology and expressed WT LMNA or E145K LMNA mutants in these cells by transient transfection (Figure S2G). Interphase cells expressing E145K LMNA mutant show a significant reduction in centromeric CENP-A, as well as CENP-C, and CENP-T compared to *LMNA KO* cells expressing WT LMNA (Figure 2A, 2C, 2D, 2F, 2G & 2I). Furthermore, the expression of the LMNA E145K mutant induced a “flowerlike” phenotypic pattern of the nuclear envelope along with the clustering of the centromeres as has been described by prior research [12]. Metaphase cells expressing E145K LMNA mutation also demonstrate a significant reduction in CENP-A, CENP-C, and CENP-T compared to *LMNA KO* cells expressing WT LMNA (Figure 2B, 2C, 2E, 2F, 2H & 2I). To study this phenotype in another system, we obtained immortalized fibroblast cells isolated from *LMNA KO* mice, and immunostained for CCAN components. Similar to HeLa *LMNA KO*, we also found a reduction in centromeric levels of CENP-C and CENP-T during both interphase and metaphase (Figure 2J-2O). Interestingly, when we monitored the CENP-T centromeric localization over time in *LMNA KO* MEF cultures, we observed that the fluorescence levels continue to progressively decrease over a period of 15 weeks (Figure S2H).

**Figure 2.**
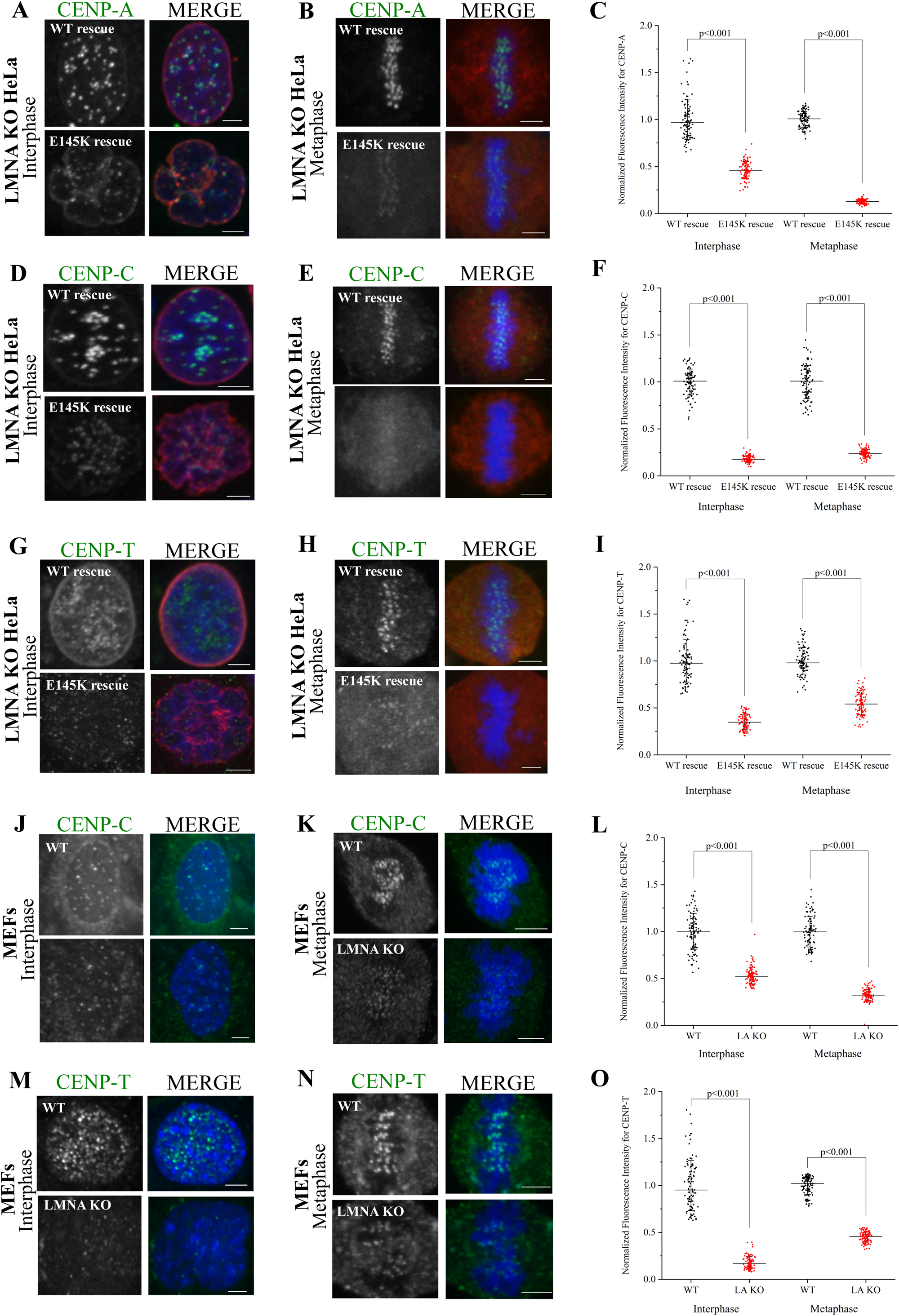
LMNA is required for proper centromere assembly in interphase and mitotic cells. (A-B) *LMNA KO* interphase (A) or mitotic (B) HeLa cells rescued with either WT or the E145K mutant of LMNA were immunostained for CENP-A (green) and the expressed FLAG-tagged LMNA (red) with the chromosomes counterstained using DAPI (blue). (C) Quantification of CENP-A immunofluorescence in the two conditions depicted in A and B. (D-E). Same as the conditions in A and B but in this case, CENP-C was detected instead of CENP-A. (F) Quantification of CENP-C immunofluorescence from D and E. (G-H). Same as the conditions in A and B, but in this case, CENP-T was detected instead of CENP-A. (I) Quantification of CENP-T immunofluorescence from G and H. (J-K) *WT* or *LMNA* (also labelled as *LA*) *KO* interphase (J) or mitotic (K) Mouse Embryonic Fibroblasts (MEFs) were immunostained for CENP-C (green) with the chromosomes counterstained using DAPI. (L) Quantification of CENP-C immunofluorescence in the two conditions depicted in J and K. (M-N). Same as the conditions in J and K, but in this case, CENP-T was detected instead of CENP-C. (O) Quantification of CENP-C immunofluorescence in the two conditions depicted in M and N. Scale bars, 5 µm.

To test whether the loss of kinetochore levels of these proteins is brought about by changes in gene expression after inhibiting LMNA function, we carried out RNA-seq experiments of control and *LMNA KO* MEFs. Our analysis revealed that the expression of none of the kinetochore genes assessed were altered after the knockout (*KO*) of *LMNA* (Figure S2I). Together, these results support the finding that inhibition of LMNA function or the expression of mutant LMNA reduces the localization of centromeric CENP-A and the CCAN components in interphase and mitotic cells, suggesting that LMNA is critical in the assembly of these centromeric components.

### Knockdown of LMNA-associated proteins, BANF1 and EMD, causes defective centromere assembly

To build upon the results of the LMNA knockdown and rescue experiments, we focused on LMNA-associated proteins (LAAPs) that are known to bind to and form a complex with LMNA/C. It is also known that BANF1 and EMD associate with the nuclear lamina underlining the nuclear envelope and that they colocalize at the core domain of chromosomes during nuclear envelope reformation [13, 14]. To determine whether inhibiting these LAAPs impact CCAN proteins in HeLa cells, we tested if CENP-A, CENP-C, and CENP-T levels at the centromeres were affected after the depletion of BANF1 by siRNA (siBANF1) [16]. Similar to that of *LMNA KO*, Western blotting of siBANF1 cell lysates showed no reduction in the cellular levels of CENP-A or CENP-C (Figure S2J). In control interphase cells, we find bright CENP-A, CENP-C, and CENP-T foci scattered throughout the nucleus as expected. However, in siBANF1 cells, we find that there was a reduction in fluorescence intensity of the foci of all the three proteins within the interphase nuclei as compared to those of the control cells (Figures S3A, S3C-top, S3D, S3F-top, S3G, and S3I). Similar to interphase cells, mitotic cells in metaphase show a significant reduction in fluorescence signal of these proteins after the perturbation of BANF1 (Figures S3B, S3C, S3E, S3F, S3H & S3I-top). Furthermore, we found that perturbations of LAAPs impacted the outer kinetochore assembly as well, as expected. For example, our data showed that siBANF1 impacted BUB1 protein levels considerably during prometaphase (Figure S4A and S4C). BUB1 is a protein kinase that plays a role in spindle assembly checkpoint and contributes to kinetochore-microtubule attachment stability [5, 7]. Additionally, we also found that *LMNA KO* MEFs showed an even greater reduction in BUB1 signal after being maintained in culture for 48-72 hours (Figures S4B & S4C).

To further demonstrate that LAAP perturbation affects centromere and the inner kinetochore, we depleted HeLa cells of another LAAP, EMD, by siRNA treatment (siEMD). Again, we found that CENP-A, CENP-C, and CENP-T fluorescent signals were significantly impacted, as the centromeric levels of these proteins were substantially reduced following siEMD in interphase as well as metaphase cells (Figure S3A-3I). However, the overall reduction of siEMD was not as potent as siBANF1 HeLa cells. To determine whether the outer kinetochore is impacted upon siEMD, we tested if the localization of ZWINT-1, an outer kinetochore protein involved in kinetochore-microtubule attachments and the recruitment of SAC proteins [5, 7], was affected after the depletion of EMD. We found that the knockdown of EMD in HeLa and RPE1 cells, both showed a significant reduction in ZWINT-1 (Figure S4D-4F). Together these results further support the idea that the localization of LAAPs to the KPC is linked to centromeric/kinetochore protein assembly. As observed with siBANF1 and *LMNA KO*, Western blotting analysis of siEMD cell lysates also showed no reduction in the cellular levels of CENP-A and CENP-C (Figure S2I).

### Inhibition of LAAP function interferes with proper mitotic progression and induces chromosome mis-segregation

Our data so far conclusively suggests that there is a functional link between LMNA-associated proteins and centromeric components that is important for kinetochore assembly. Previous studies show defects in chromosome segregation after the perturbation of LAAP genes [17, 18]. To determine whether kinetochore function and mitotic progression occur properly when LAAPs are perturbed, we analyzed mitotic *LMNA KO* cells and found that there was a considerable increase in the frequency of mitotic cells. This phenotype was manifested with a reproducible increase in the frequency of early mitotic cells in prometaphase (Figure S4G). While these results points to a delay in mitotic cells undergoing chromosome segregation, most cells progressed to anaphase and telophase. We thus analyzed chromosome segregation in anaphase and telophase, and found a greater than 3-fold increase in chromosome mis-segregation events consistent with the fact that kinetochore function was affected specifically in cells lacking LMNA (Figure S4H). To further support that LAAP inhibition affects mitotic progression and kinetochore function, we carried out a similar analysis in BANF1- or EMD-depleted cells. Similar to that of *LMNA KO*, we found that there was a consistent increase in the number of mitotic cells in prometaphase after siBANF1 (Figure S4I) or siEMD (Figure S4K). However, in the case of siBANF1, we also observed a moderate increase in the frequency of telophase cells (Figure S4I). Our current data cannot completely explain this interesting difference and further studies are required to delineate the details in this context. As with LMNA inhibition, we found that siBANF1 or siEMD also resulted in a significant increase in chromosome mis-segregation events suggesting a kinetochore dysfunction (Figure S4J and S4L). The data demonstrate that mitotic progression is abnormal after perturbing LAAP function even though the cells did not arrest prior to anaphase.

### LAAPs and centromeric components bind to each other and are co-dependent for their assembly at the kinetochore proximal core domain

Our work so far suggests that LAAPs are important for kinetochore assembly and function. As has been described, LAAP inhibition interferes with the localization of centromeric CCAN components at the KPC domain. To test if there was an inter-dependence between the LAAPs and the inner kinetochore for their recruitment to the KPC or the core domains in general, we depleted CENP-C in HeLa cells via siRNA treatment (siCENP-C) to disrupt the inner kinetochore structure. Very interestingly, we observed that there was a substantial reduction and/or mislocalization of BANF1 (Figure 3A, 3B) and EMD (Figure 3C, 3D) at both the KPC and the KDC in this scenario. We next used a Dox inducible *CENP-W* (another CCAN component) *KO* cell line to test this further. Similar to CENP-C knockdown, *CENP-W KO* resulted in defective inner kinetochore assembly as the localization of CENP-T to kinetochores was lost in this scenario (Figure 3E). Our data show that there was a severe loss of EMD from both the core domains in response to inner kinetochore assembly perturbation, suggesting that LAAPs and the CCAN are mutually dependent for their assembly at the core (Figure 3E, 3F).

**Figure 3.**
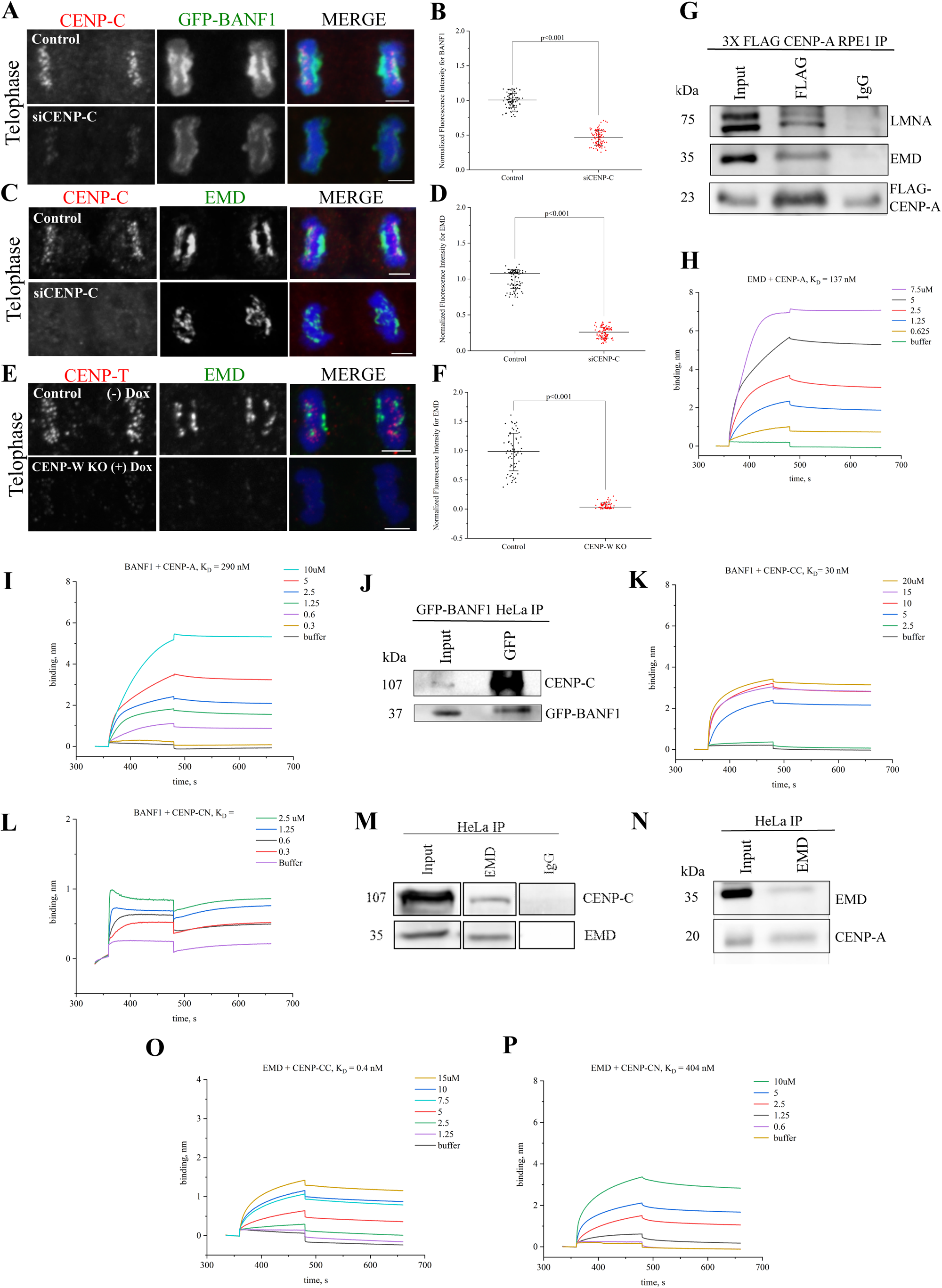
Biochemical interaction between LAAPs and the centromeric proteins facilitate their codependent assembly at the KPC. (A) HeLa cells in Mitotic telophase were immunostained for CENP-C (red) and GFP-BANF1 (green) in control cells or those treated with siRNA against CENP-C (siCENP-C) with the chromosomes counterstained using DAPI (blue). (B) Quantification of BANF1 immunofluorescence in WT or siCENP*-*C cells as depicted in A. (C) Similar to the conditions as in A, but cells were immunostained for EMD (green) instead of GFP-BANF1. (D) Quantification of EMD immunofluorescence in control or siCENP*-*C cells as depicted in C. (E) Telophase cells immunostained for CENP-T (red) and EMD (green) in WT and inducible *CENP-W KO* HeLa cells. (F) Quantification of EMD immunofluorescence in control or *CENP-W KO* cells as depicted in E. (G) RPE1 cells over-expressing FLAG-tagged CENP-A were immunoprecipitated using FLAG antibody and probed for LMNA, EMD, and FLAG. IgG was used as negative control. (H) Biolayer interferometry (BLI) sensograms obtained using EMD as the ligand with various concentrations (0.625-7.5 µM) of CENP-A as the analyte. Buffer only sample was used to set the baseline. The graph depicts the binding and dissociation phases along with equilibrium dissociation constant (K_D_) that was determined. (I) BLI sensograms obtained using BANF1 as the ligand with various concentrations (0.3-10 µM) of CENP-A as the analyte. (J) GFP-BANF1 HeLa cell lysates were immunoprecipitated with anti-GFP antibody and probed for CENP-C. (K) BLI sensogram of BANF1 with various concentrations (2.5-20 µM) of the C-terminal of CENP-C. (L) BLI sensogram of BANF1 with various concentrations (0.3-2.5 µM) of the N-terminal region of CENP-C. (M-N) Immunoprecipitation of HeLa cell lysates showing EMD pulling down both CENP-C (M) as well as CENP-A (N). (O) BLI sensogram of EMD with various concentrations (1.25-15 µM) of the C-terminal region of CENP-C. (P) BLI sensogram of EMD with various concentrations (0.6-10 µM) of the N-terminal region of CENP-C. Scale bars, 5 µm.

After demonstrating a critical structural and functional link between LAAPs and the centromeres, we now wanted to obtain biochemical evidence for a direct link between these components. We thus decided to test for an interaction between the centromeric and LAAP components using RPE1 cells overexpressing FLAG-tagged CENP-A. Based on our findings so far, we reasoned that the critical interaction between the centromeres and LAAP components possibly occurred in telophase and/or early G1 cells. We thus arrested cells in G2/M transition by treating them with the CDK1 inhibitor, RO-3306, for 19 hours and prepared cell extracts 1.5 hours after release from the arrest to obtain maximum cells in telophase. The cell lysates were subjected to co-immunoprecipitation (Co-IP) using antibodies against FLAG-tag and the immunoprecipitates were probed with antibodies against EMD and LMNA after transfer to membranes. Strikingly we observed that both LMNA as well as EMD were successfully immunoprecipitated with CENP-A while a control IgG antibody immunoprecipitated neither of these components (Figure 3G). Having assessed this interaction *in vivo,* we next sought to confirm it by determining the affinity of binding between EMD and CENP-A using *in vitro* Bio-layer interferometry (BLI). For this experiment, EMD was the immobilized ligand while CENP-A was used as the analyte in solution. Based on our results, we observed that EMD binds to CENP-A with a high affinity of 137 nM measured K_D_. (Figure 3H). Previous studies had demonstrated that BANF1 can bind to Histone H3 [19], of which centromeric CENP-A is a variant. We thus tested if we could also observe a direct interaction between BANF1 and CENP-A using the same BLI assay where purified BANF1 was used as a ligand and CENP-A was the analyte. We observed that these two proteins also interacted with each other *in vitro*, but in this case the affinity (K_D_ = 290 nM) was slightly lower as compared to EMD-CENP-A binding (Figure 3I).

As studies in *Drosophila* S2 cells have shown an interaction between BANF1 and CENP-C [10], we next tested if CENP-C can also interact with human BANF1. We first performed Co-IP experiments in HeLa cells stably expressing GFP-BANF1 employing our RO-3306 arrest and release protocol. The cell lysates were subjected to immunoprecipitation using an antibody against GFP and immunoblotted using an anti-CENP-C antibody. As expected, BANF1 indeed immunoprecipitated CENP-C (Figure 3J). Having observed an interaction *in vivo,* we next sought to investigate the extent of the binding between BANF1 and CENP-C using BLI. BANF1 was the immobilized ligand while either purified CENP-C N-terminal or C-terminal region was used as the analyte. Based on our results, BANF1 bound preferentially to the C-terminus of CENP-C as determined by its equilibrium constant K_D_ which was 30nM (Figure 3K) while BANF1 did not show any binding to the N-terminus of CENP-C, the K_D_ for which could thus not be determined (Figure 3L).

Based on our findings so far, we also tested for a possible interaction between EMD and CENP-C using both immunoprecipitation and BLI experiments. An antibody against EMD was used for IPs of HeLa cell lysates obtained from telophase-synchronized cells. EMD immunoprecipitated both CENP-C and CENP-A (Figure 3M and 3N). Next, we tested EMD’s binding affinity for the N-terminus and/or C-terminus of CENP-C using BLI. EMD in this case was used as a ligand while either of the two CENP-C protein fragments were used as analytes. We determined that similar to BANF1, EMD also had a much higher binding affinity for the C-terminus of CENP-C as its K_D_ was shown to be 0.4 nM (Figure 3O), while the K_D_ for binding the CENP-C N-terminus was determined to be 404 nM (Figure 3P). Together this data conclusively confirms multiple modes of interaction between the LAAPs and the centromere/kinetochore components, where EMD seems to serve a central function.

### The localization of factors that load new CENP-A to the centromeres during M-G1 cell cycle transition is compromised in LAAP-inhibited cells

Having identified the essential biochemical links that connect centromeres to the core domain, we next intended to obtain mechanistic insights into the relevance of this connection. The key structural component that defines the centromere, CENP-A, is loaded very late in telophase and in the succeeding G1 phase of the cell cycle. The timing of new CENP-A loading has been shown to be initiated at ∼50 mins after anaphase onset and continues for several hours in the G1 phase [20]. On the other hand, the timing of the loading of the LAAP components including BANF1, EMD and LMNA has been mapped to very early stages of telophase within minutes after anaphase onset. BANF1 is loaded as early as 6-7 mins after anaphase onset (Figure S1C; Video S1), followed by EMD immediately after that and LMNA in 9-10 mins (Figure 1F; Video S2) [13–15]. Loading of new CENP-A onto the centromeres requires the Mis18 adapter complex and the HJURP histone chaperone [21–25]. Mis18 loading is the first step in the process and is in turn required for the recruitment and regulation of HJURP-mediated CENP-A deposition at the centromeric chromatin [26, 27]. Interestingly, while HJURP is functional in CENP-A loading only in G1 and S phases after a mitotic cycle, the Mis18 complex is loaded in telophase as early as between 10-20 mins after anaphase onset, which is immediately after the point when LAAP loading is known to occur [22]. Consistent with that, we observed a striking co-distribution of Mis18 components, GFP-Mis18α and Mis18BP1 with EMD in telophase (Figure 4A, top panel).

**Figure 4.**
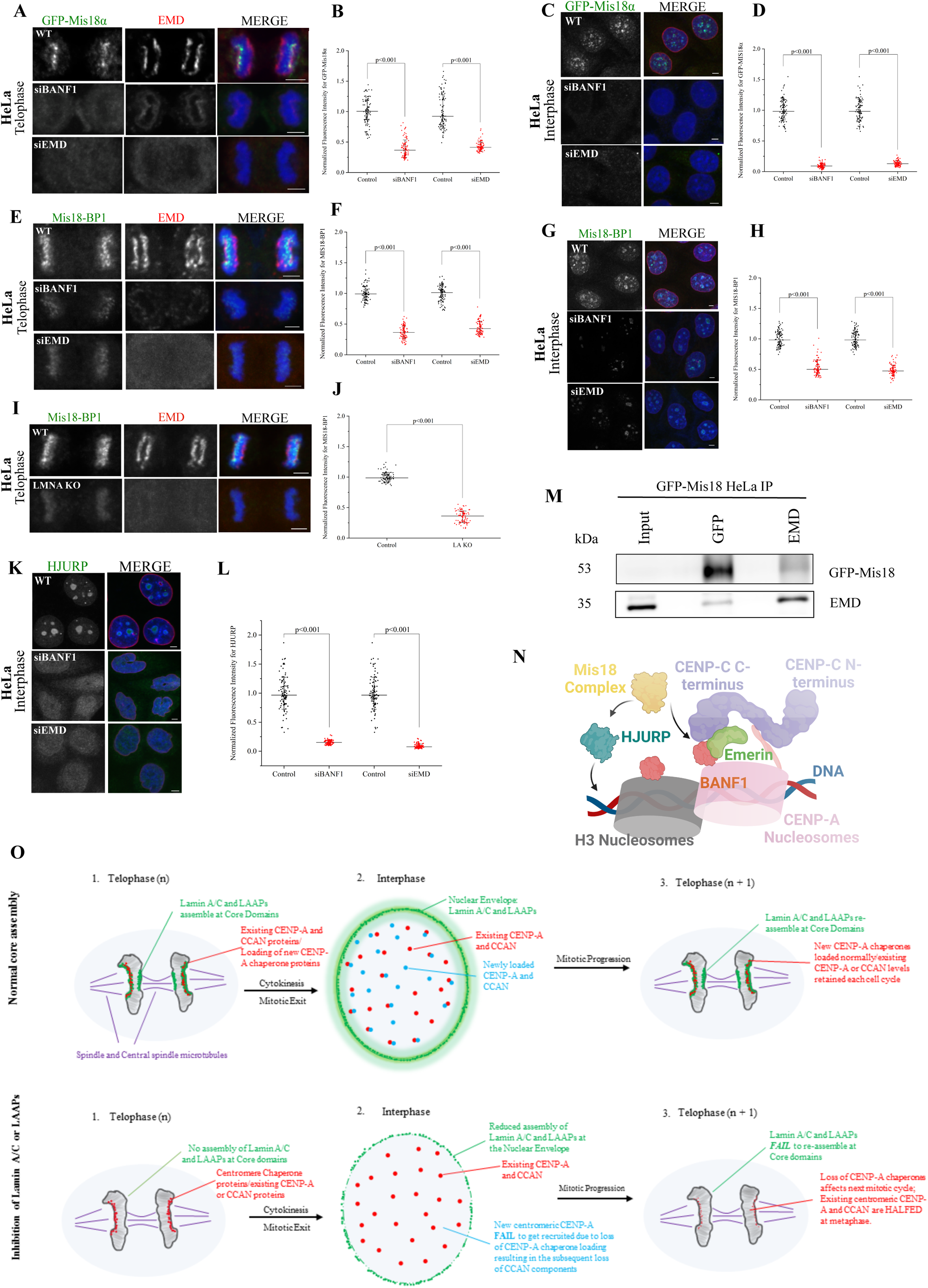
The loading of CENP-A adaptor Mis18 and the CENP-A chaperone, HJURP, to the centromeres is severely affected in LAAP-inhibited cells. (A) Control, siBANF1- or siEMD-treated mitotic telophase HeLa cells constitutively expressing the GFP-Mis18α subunit of the Mis18 complex were immunostained for GFP-Mis18α (green) and EMD (red) with chromosomes counterstained using DAPI (blue). (B) Quantification of GFP-Mis18α immunofluorescence in the conditions as depicted in A. (C) same conditions as in A but with interphase cells. (D) Quantification of GFP-Mis18α immunofluorescence in the conditions as depicted in C. (E) Control, siBANF1 or siEMD mitotic telophase HeLa cells were immunostained for the Mis18 complex subunit, Mis18-BP1 (green), and EMD (red) with chromosomes counterstained using DAPI. (F) Quantification of Mis18-BP1 immunofluorescence in the conditions as depicted in E. (G) same conditions as in E but with interphase cells. (H) Quantification of Mis18-BP1 immunofluorescence in the conditions as depicted in G. (I) *WT* and *LMNA KO* HeLa cells in mitotic telophase were immunostained for Mis18-BP1 (green) and EMD (red). (J) Quantification of Mis18-BP1 immunofluorescence in the conditions as depicted in I. (K) Control, siBANF1, and siEMD HeLa cells in interphase were immunostained for HJURP (green) with the chromosomes counterstained using DAPI. (L) Quantification of HJURP immunofluorescence in the conditions as depicted in K. Scale bars, 5 µm. (M) Immunoprecipitation of GFP-Mis18α HeLa cell lysates using anti-GFP and anti-EMD antibodies with the blot being reversibly probed for both these proteins. (N) A Molecular Model for LAAP-centromere co-assembly. The centromeric DNA and the CCAN is instrumental in the recruitment of LAAPs during telophase. Once the core is assembled, both these structures (CCAN and the core) seems to contribute to the loading of the Mis18 complex later in telophase. Mis18 along with HJURP can now recruit new copies of CENP-A in late telophase and G1. (O) A Schematic model of core function in normal cell division (top) and cells in which LAAPs are perturbed (bottom). During normal core assembly, LMNA and the LAAPs (green) assemble at the two core domains during telophase, contributing to the recruitment of CENP-A chaperones (red). The existing centromeric and CCAN components contribute to the assembly of LAAPs at the core domains. Following cytokinesis and entry into the next G1, LMNA and the LAAPs relocate from the cores to the nuclear envelope while new CENP-A as well as new CCANs components (yellow) gets loaded onto the centromeres so that there is adequate amounts of CENP-A and CCAN components retained for the succeeding round of mitosis. However, when LMNA or LAAPs are perturbed, these proteins obviously do not assemble at the core domains during telophase. This results in a substantial reduction of CENP-A chaperone recruitment in telophase, which in turn interferes with new CENP-A and CCAN recruitment to centromeres. When cells progress through multiple rounds of cell division cycle, without LMNA or LAAPs assembly at the core, there is a continuous loss of CENP-A and CCANs proteins from the centromeres, leading to the manifestation of severe kinetochore assembly defects and rampant chromosome mis-segregation.

Based on these observations, we tested the dependence of new Mis18 loading in telophase on the deposition of LAAPs at the KPC domain. We used HeLa cells constitutively expressing GFP-Mis18α subunit of the Mis18 complex and the same inhibition approaches used to perturb the core previously, including *LMNA KO*/rescue or the knockdown of BANF1/EMD. Strikingly, we observed that independent of the approach used, there was a severe decrease in the levels of new Mis18α subunits loaded in telophase after the perturbation of LAAP function. Both BANF1 as well as EMD depletion resulted in substantial loss of Mis18α from telophase centromeres (Figure 4A, 4B). To test for Mis18 localization to centromeric foci within the interphase nucleus, we combined these inhibition protocols with a cell synchronization approach involving R03306 treatment/release followed by MG132 treatment/release so that we obtained a preponderance of cells in early G1. Consistent with the results obtained in telophase, the fluorescence levels of Mis18α centromeric foci was substantially reduced (Figure 4C, 4D). Further, another subunit of the Mis18 complex, Mis18BP1, also follow an identical trend in that the localization of this component to centromeric loci in telophase and in interphase cells were substantially reduced in BANF1 or EMD depleted cells (Figure 4E-4H) as well as in *LMNA KO* cells (Figure 4I, 4J). As the loading of the CENP-A chaperone HJURP within the interphase nuclei is dependent on Mis18 [26, 27], we next tested if this process was also perturbed by the inhibition of LAAP-localization to the KPC. Indeed, we observed a similar trend with the centromeric HJURP localization pattern in interphase cells (Figure 4K-4L). Finally, we found that GFP-Mis18α as well as EMD pulled each other down from HeLa cell lysates while using anti-GFP and anti-EMD antibodies respectively in immunoprecipitation experiments (Figure 4M), suggesting that LAAP deposition at the core domain could be directly connected to the Mis18 loading in telophase.

It has been previously demonstrated that the deleterious effects of the LMNA E145K mutations were primarily post-mitotic [12] in that the cells possibly need to go through mitotic divisions over multiple cell cycles prior to the manifestation of the phenotype (Figure S2H). It has also been shown that it takes several rounds of cell cycles for the severe effects of CENP-A loss to manifest with regard to chromosome segregation and mitotic progression [28]. We predict that human cells establish a variant steady-state where progressively reducing levels of centromeric and kinetochore components after LAAP perturbation is still competent to perform chromosome alignment and segregation (albeit with errors in the process), prior to their mitotic progression being completely blocked and triggering cell death.

Our findings in this study leads to a model where the co-deposition of the LAAP with the centromeric components are critical for the loading of the CENP-A chaperones in telophase and early G1 cells and is hence critical for the subsequent loading of new centromeric and CCAN components in the succeeding interphase (Model in Figure 4O). An important outstanding question is if the assembly of the LAAP proteins at the kinetochore-distal core (KDC) domain where there are no centromeres present, is still important for centromere assembly. It is not clear at this point if this assembly could still be important somehow for the loading of the Mis18 complex and consequently for the assembly of new centromeric components. As preventing centromere assembly affects the deposition of LAAPs at both KPC and KDC domains, our current results support a possibility that the initial deposition of the LAAPs at the KPC might in turn control further LAAP localization at the KDC at a slightly later time point by diffusion or other unknown mechanisms. Alternatively, since BANF1 also binds to Histone H3 [19], it is possible that the presence of CENP-A nucleosomes proximal to the KPC is inconsequential to LAAP recruitment and that LAAPs are recruited to both the KPC and the KDC simultaneously. The precise nature of dependencies for Mis18 loading in telophase between the mechanisms mediated by CENP-C [29, 30] and that mediated by the LAAP is also unclear at this point. Time-resolved live-imaging as well as biochemical studies in the future should address these outstanding questions in our efforts to better define the enigmatic connection between nuclear envelope reformation and centromere assembly.

## Materials and Methods

### Cell Culture and Drug Treatments

HeLa cells or those stably expressing human proteins (including *LMNA KO*, *CENP-W KO*, GFP-BANF1, GFP-Mis18α), MEFs, and RPE1 cells were cultured in complete Dulbecco’s modified Eagle’s medium (DMEM; Corning #10-013-CV) supplemented with 10% fetal bovine serum (Gemini Bio, # 900-308-500) and Penicillin-streptomycin (100 U ml^−1^ penicillin and 1μg ml^−1^ streptomycin) at 37°C and 5% CO_2_. *LMNA KO* MEFs was provided by Robert Goldman lab at Northwestern University [31]; GFP-BANF1 HeLa cell line was provided by Daniel Gerlich lab at IMBA, Vienna, Austria [16]; *CENP-W* KO HeLa cell lines were provided by Iain Cheeseman’s lab at MIT [32]; GFP-Mis18α stable HeLa was provided by Dan Foltz’s lab at Northwestern University [26]. Synchronization of cells in telophase or early G1 was achieved through drug treatments. Cells were arrested at G2/M using RO-3306 treatment (9 μM) for 19 hours. Cells were then released from drug treatment and allowed to proceed to telophase for 1.5 hours. For synchronizing in early G1, RO-3306 treatment/release was combined with MG132 treatment/release. Cells were released into 10 μM MG132 for 2 hours after RO-3306 treatment. Maximum synchrony in G1 was obtained after 45 min-1 hour of release from MG132.

### siRNA Knockdown

Depletion of BANF1, EMD, LMNA and CENP-C were achieved through siRNA-mediated knockdown. siRNAs were synthesized either as separate oligonucleotides (Invitrogen-ThermoFisher Inc.) or as a SMARTpool of multiple oligonucleotides (Dharmacon - GE Healthcare) and transfected using Lipofectamine RNAiMax as per manufacturer’s instructions (Invitrogen -ThermoFisher Inc). Cells were seeded onto flame-sterilized 18×18mm square glass coverslips (CORNING, #2850-18) in 6-well plates. For BANF1 knockdown, reverse transfection protocols were followed in which siBANF1 was transfected at the time of cell seeding as described in Samwer et al., 2017 [16]. The siRNA target sequence used for BANF1 was AGUUUCUGGUGCUAAAGAA[DT][DT] (Invitrogen-ThermoFisher Inc.) For all other knockdown approaches, forward transfection protocols were followed in which siRNA(s), single or multiple, was transfected 24 hours after cell seeding. For EMD, a mixture of three siEMD oligonucleotide target sequences: CCA GGU GCG UGA UGA CAU U[d T][dT], GAG CAA GGA CUA UGA U[dT][dT], CUU UGU UUA CUA UUC CAU A[dT][dT] were used (Invitrogen - ThermoFisher Inc.)[33]. For LMNA, we used LMNA siRNA - ONTARGET plus SMARTpool, L004978-00 (Dharmacon - GE Healthcare). For CENP-C, we used CENP-C siRNA - ONTARGET plus SMARTpool, L003251-00-0005 (Dharmacon - Horizon Discovery Group). Cells were fixed onto coverslips 72 hours (for CENP-C, LMNA and EMD knockdown) or 96 hours (for BANF1 knockdown) post-transfection.

### Generating HeLa LMNA KO cells

The gRNAs targeting the human *LMNA* gene (NCBI-Gene ID: 4000) in HeLa cells were designed using CRISPR based Gene Knockout kit v2 (Synthego Corporation). The sequences of different multiguide RNAs were: C*U*U*UAGCAAUACCAAGAAGG, G*G*C*UCUGCUGAACUCCAAGG, G*C*A*UGAUCUGCGGGGCCAGG. HeLa cells were seeded and propagated to desired confluency in a 37°C incubator with 5% CO_2_. The *LMNA* gRNAs were incubated with spCas9 2NLS protein for 20 min at RT to make the complex of Cas9 RNPs. The RNP complex was then electroporated into HeLa cells using the Neon NxT Electroporation System (ThermoFisher Inc.) under the kit conditions provided by the manufacturer, Synthego. For positive control, the same reaction was performed with a Synthego control gRNA, which was not specific to the human genome. The electroporated cells were cultured for 24-48 hours and sorted as singles cells into 96-well plates containing DMEM medium supplemented with 20% FBS. We then monitored the cell growth of single colonies for 4-6 weeks while the cells grew into colonies, followed by scaling up into 24-well, and then into 6-well plates. The cells were then split into two groups, one for cryopreservation and the other to validate by Western blot to confirm homozygous knockout clones.

### LMNA mutant Rescue Experiments

WT and E145K mutant rescue experiments employing *LMNA KO* HeLa cells were achieved by transient transfection using Effectene transfection reagent (Qiagen, # 301425). pCMV2 plasmids expressing FLAG-tagged WT and E145K LMNA constructs (a gift from Dr. Robert Goldman lab, Northwestern University) were transfected 24 hours after seeding of cells on coverslips in 6-well plates using a working concentration of 0.25-0.4 ug/ul DNA at a 1:25 DNA:Effectene ratio. Cells were fixed in PFA or Methanol (details below) 48 hours following transfection. The coverslips were stained with either anti-FLAG mouse monoclonal or rabbit polyclonal antibodies depending on the other antibodies used for co-staining.

### RNA Sequencing

RNA was prepared from *WT* and *LMNA KO* MEFs using the Direct-zol^TM^ RNA MiniPrep kit from Zymo Research (#R2050). RNA quality control was performed at the NUSeq Core facility at Northwestern University. Library preparation, sequencing and data analysis (Standard Analysis Package) was performed using the Standard RNA-seq services from GENEWIZ, Inc. The Volcano plot depiction of the analyzed data was generated using the R Program.

### Immunofluorescence microscopy and antibodies

Cells were fixed onto coverslips either with 100% ice-cold methanol or 3.6% Paraformaldehyde. Coverslips were first rinsed three times in 1x PBS. For methanol fixation, cells were pre-fixed with ice-cold methanol for 30 sec-1 min, then incubated at −20℃ for 5-6 mins. Alternatively, cells were permeabilized with 0.5% Triton X-100 for 5 mins, then fixed with 3.6% Paraformaldehyde for 20 mins at room temperature (RT). Blocking was performed with 0.1% BSA in PBS for 1 hour at RT. Cells were immunostained with primary antibody overnight at 4℃, washed three times with 1x PBS for 10 mins each, then incubated with Alexa Fluor-488/647 or Rhodamine Red-X (Thermo Fisher Scientific) secondary antibody for 1 hour at 37℃. After two more washes in PBS, nuclei/chromosomes were counterstained with DAPI in 1x PBS (1:10,000) for 10 mins at RT and the coverslips were mounted onto glass slides with ProLong Gold Antifade reagent (Thermo Fisher Scientific) mounting media. Slides were stored at 4℃ until visualization.

The antibodies used for immunofluorescence include anti-human ACA (Immunovision Inc., #HCT-0100), anti-HJURP rabbit polyclonal (a kind gift from Dr. Dan Foltz, Northwestern University), anti-LMNA mouse and rabbit antibodies (kind gifts from Dr. Robert Goldman, Northwestern University), anti-CENP-A mouse monoclonal (Invitrogen-ThermoFisher Inc 3-19, #MA1-20832), anti-CENP-C mouse monoclonal (abcam, #ab50974), anti-CENP-C rabbit monoclonal (abcam, #ab193666), anti-CENP-T rabbit polyclonal (abcam, #ab220280), anti-ZWINT-1 rabbit polyclonal (Bethyl Laboratory Inc., #IHC-00095), anti-Bub1 mouse monoclonal (abcam, #ab54893), anti-FLAG rabbit polyclonal (Sigma-Aldrich, #F7425), anti-FLAG M2 mouse monoclonal (Sigma-Aldrich, #F3165), anti-EMD rabbit polyclonal (Proteintech, # 10351-1-AP), anti-EMD mouse monoclonal (Novocastra, #NCL-EMERIN), anti-GFP rabbit polyclonal (Invitrogen, #A11122), and anti-Mis18BP1 rabbit polyclonal antibody (a kind gift from Dr. Paul Maddox, UNC-Chapel Hill).

### Image acquisition and Processing

For fixed-cell imaging, three-dimensional stacks were obtained through the cells using a Nikon Eclipse TiE inverted microscope equipped with a Yokogawa CSU-X1 spinning disc, an Andor iXon Ultra888 EMCCD camera and an x60 or x100 1.4 NA Plan-Apochromatic DIC oil immersion objective (Nikon). For fixed cell experiments, images were acquired at RT as Z-stacks at 0.2-0.3 µm intervals controlled by NIS-elements software (Nikon). Images were processed in NIS-Elements software (Nikon) and represented as maximum-intensity projections of the required z-stacks. A total of 100 centromeric foci from at least 5 different interphase or mitotic cells were analyzed for the fluorescence quantifications.

For 3D-SIM super-resolution imaging, cells were imaged on a Nikon N-SIM system using a Nikon Plan Apo 100×1.49NA TIRF objective. The fluorophores used were Alexa488 and Rhodaomine Red-X; Prolong Gold Antifade mounting media was used to mount the coverslips. Images were captured using Nikon Elements software with a Hamamatsu Flash 4 camera. Z-step size was set to 0.100 µm (well within Nyquist criterion). 15 images per slice (5 phases, 3 angles) were captured and then reconstructed also using the Nikon Elements software.

### Live cell imaging

Live imaging was carried out on 35-mm glass-bottomed dishes (MatTek Corporation) in an incubation chamber for microscopes (Tokai Hit Co., Ltd) at 37°C and 5% CO_2_ using FluoroBrite DMEM live imaging media (Life Technologies) supplemented with 2.5% FBS. For the live imaging of WT and E145K mutant LMNA, HeLa cells stably expressing Histone H2B-RFP were transfected with the respective GFP-tagged LMNA variant constructs for 48 hours before initiating live imaging. For the live imaging of GFP-BANF1 stable HeLa cells, the chromosomes were labeled using the membrane permeable DNA stain, Hoechst 33258 for 1.5 hours before initiating live imaging. Images were acquired every 30 sec (for GFP-BANF1) or every 1 min (for GFP-LMNA) for up to 25-30 mins as required.

### Co-Immunoprecipitation and Western Blotting

For the preparation of whole cell lysates, cells were lysed with RIPA buffer (Sigma-Aldrich, #R0278) containing Halt Protease Inhibitor Cocktail (Thermo Scientific, #87786) and incubated on ice for 20 minutes. Samples were centrifuged at 14k rpm at 4℃ for 5 minutes and the supernatant was collected. Protein concentrations were determined with Coomassie protein assay, then samples were prepared with Laemmli Sample Buffer and boiled at 100℃ for 5 minutes. Samples were resolved on 12-15% SDS-PAGE gel, then transferred to PVDF blotting membrane (Cytiva Amersham Hybond, #10600023).

For Western blotting, PVDF membrane was blocked in 5% non-fat dry milk made in 1x TBS with 0.1% Tween20 solution (TBST) for 1 hour at RT with shaking, followed by three 10 minute washes with 1x TBST. Primary and secondary antibodies were prepared in 1x TBST with 5% BSA. The membranes were incubated at RT shaking with appropriate primary antibodies for 1 hour, washed three times with 1x TBST, then incubated with goat anti-rabbit (Azure Biosystems, #AC2114) or goat anti-mouse HRP secondary antibodies (Azure Biosystems, #AC2115) used at 1:2000 for 1 hour shaking at RT. Detection was achieved using SuperSignal West Pico PLUS Chemiluminescent Substrate (Thermo Fisher Scientific, #34580).

Primary antibodies used for Western blot include anti-α-Actin (1A4) mouse monoclonal (Santa Cruz, #sc-32251), beta Actin (C4) mouse monoclonal (Santa Cruz, #sc-47778), anti-EMD rabbit polyclonal (Proteintech, #10351-1-AP), anti-LMNA/C rabbit polyclonal and mouse monoclonal (a kind gift from Dr. Robert Goldman lab, Northwestern University), anti-CENP-C rabbit monoclonal (Abcam, #ab193666), anti-CENP-A mouse monoclonal 3-19 (Invitrogen, # MA1-20832), anti-FLAG M2 mouse monoclonal (Sigma-Aldrich #F3165), and anti-GFP rabbit polyclonal (Invitrogen, #A11122).

For Co-immunoprecipitation, cells were lysed using Pierce IP Lysis Buffer (Thermo Scientific, #87787) with Halt Protease Inhibitor Cocktail. Protein concentration was determined using Coomassie protein assay (Thermo Scientific, #23200). Samples were precleared, then incubated overnight rotating at 4℃ with antibody, followed by 2 hour incubation at 4℃ with 50 µL of magnetic Dynabeads Protein G (Invitrogen, #10004D). Beads were washed with 1x TBS Tween-20 washing buffer (Thermo Scientific, #28360) then separated from the protein by boiling for 10 mins at 100℃ with 4x Laemmli Sample Buffer (BioRad, #1610747).

The antibodies used for immunoprecipitation include anti-GFP rabbit polyclonal (Invitrogen, #A11122), anti-EMD rabbit polyclonal (Proteintech #10351-1-AP), and anti-FLAG M2 mouse monoclonal (Sigma-Aldrich #F3165).

### Protein purification

Cloning and purification for Human EMD: The ORF corresponding to the Human EMD (Uni Prot. ID: NP_000108) was PCR amplified using a Human EMD plasmid cloned in pEGFP-N1 vector (a gift from Gant Luxton [34]) as a template and the indicated primers: FP-GATCGAGGAAAACCTGTACTTCCAATCCAATATGGACAACTACGCAGATCTTTCG, RP-TCGAATTCGGATCCGTTATCCACTTCCAATCTAGAAGGGGTTGCCTTCTTCAGC. The amplified DNA fragments were ligated to the pET His6 GFP TEV LIC expression vector [1GFP], Addgene plasmid 29663 using Gibson Assembly. The recombinant Human EMD protein with N-terminal His-tag was expressed in *E. coli* BL21(λDE3) by inducing with 0.25 mM IPTG. The protein was purified from soluble fraction under native conditions using Ni2+– NTA affinity chromatography (Ni2+ beads obtained from Qiagen, #30210), followed by a second step of purification involving gel filtration chromatography using Superdex 200 10/300 GL (GE Healthcare) mounted on an AKTApure system.

Purification for BANF1: For BANF1 purification, BANF1 plasmid (pBAF plasmid Addgene #104152) was transformed into *E. coli* BL21(λDE3) cells and induced with 0.5 mM IPTG for 4 h at 37 °C. The cell pellet was lysed by sonication and centrifuged at 14k for 30 mins. Since BANF1 is known to be insoluble [35], the lysed pellet obtained was resuspended in a buffer containing Guanidium Chloride (GdnCl) (Buffer B: 6M GdnCl in 50 mM HEPES and 150 mM NaCl, pH 7.5) and incubated at RT with shaking overnight. The next day, cells were centrifuged at 35k, RT, for 45 minutes. The supernatant was collected and allowed to bind with TALON® Metal Affinity Resin (Clontech Laboratories #635503) for 1 hour at RT with rotation. The supernatant was passed through a column, and the protein was refolded on the beads using a concentration gradient of GdnCl from 6 M to 0 M (Buffer A: 50 mM HEPES and 150 mM NaCl, pH 7.5). Elution was performed with imidazole and the second step of purification was performed using Superose 6 Increase 10/300 GL gel filtration chromatography (GE Healthcare).

### Bio-layer Interferometry (BLI)

Experiments were performed with a single-channel BLItz system by *forte*BIO as described in [36]. Anti-Penta-HIS (HIS1K, FORTEBIO, #185121) Biosensors were first hydrated in a black 96-well plate containing HEPES buffer (20 mM HEPES, 150 mM NaCl, and 0.1% Tween 20, pH 7.4) for 10 mins. The ligands, BANF1 or EMD, were prepared in the same buffer and immobilized to the sensor during loading phase for 300 s. Various analyte concentrations (0.3-20 uM) of 6xHis-MBP-CENP-C^1-600^-SpyCatcher, MBP-SpyTag-CENP-C^601-943^-8xHis (kind gifts from Dr, Andrea Musacchio, Max Planck Institute, Dortmund, Germany; [37], or CENP-A/H4 [1.9 mg/ml (0.3 mM) in 20mM SodPhos (pH 6.9), 2M NaCl, 14mM BME, and TRACE PMSF; a kind gift from Dr. Dan Foltz, Northwestern University] were also prepared in the same HEPES buffer. Analyte samples were loaded onto a 4.0 μl magnetic drop holder and allowed to interact with the ligand during the association phase for 120 s. The dissociation phase was set for 180 s. HEPES buffer alone was used as a baseline. All steps were performed with shaking enabled at 1000 rpm. The program BLItz Pro Software ver.1.1.0.29 was used for analysis.

### Statistical Analysis

Graphs and data plots were designed using Origin 2018 software. Statistical significance was determined by Mann-Whitney U-test.

## Acknowledgements

As detailed in the methods, we would like to thank Dr. Robert Goldman (Northwestern University) for various antibodies, cell lines and plasmids; Alexander Lee, and Justin Bodner from Dr. Daniel Foltz lab (Northwestern University) for various help with the reagents; Drs. Andrea Musacchio and Marion Pesenti for the CENP-C protein fragments; Dr. Iain Cheeseman for the *CENP-W KO* HeLa cell line; Dr. Paul Maddox for the Mis18BP1 antibody; and Dr. Daniel Gerlich for the GFP-BANF1 cell line. We would also like to acknowledge the Northwestern University Center for Advanced Microscopy for access and help with the Nikon N-SIM system, purchased through the support of NIH 1S10OD016342-01. This work was supported by NIGMS grant R01GM135391 to DV; and NIGMS grant R01GM111907 to DRF.

---

**Supplemental Figure 1. LAAPs co-localize with the centromeres at the KPC in telophase but not within the interphase nuclei.**

(A) Top panel: HeLa cells in interphase with their nuclei counterstained using DAPI (blue) were immunostained for CENP-A (red) and GFP-BANF1 (green). 2nd panel: Same as in the top panel but in this case, the cells were also immunostained for CENP-C (red) and EMD (green). 3rd panel: Same as in the top panel but in this case the cells were immunostained for CENP-A (red) and EMD (green). Fourth panel: Mouse Embryonic Fibroblast (MEFs) cells in interphase with their nuclei counterstained using DAPI (blue) were immunostained for CENP-C (red) and LMNA (green). Bottom panel: RPE1 cells in interphase with their nuclei counterstained using DAPI (blue) were immunostained for CENP-T (red) and LMNA (green). (B) Mitotic telophase HeLa cells were immunostained for EMD (green) in combination with CENP-A (top panel), CENP-C (middle panel), and CENP-T (bottom panel) and the chromosomes were counterstained using DAPI. (C) Live-cell imaging of GFP-BANF1-expressing HeLa cells at various time points after anaphase onset showing GFP-BANF1 (green) loading onto the chromosomes that were stained using Hoechst. Scale bars, 5 µm.

**Supplemental Figure 2. Inhibition of LMNA interferes with proper inner kinetochore assembly.** (A-B) Control HeLa cells (top panels) or those subjected to LMNA siRNA (siLMNA or siLA/C) treatment (bottom panels) were immunostained for CENP-C (green), and anti-CREST antiserum (ACA, red) with their chromosomes counterstained using DAPI (blue) as indicated. Interphase cells are depicted in A while late telophase cells are depicted in B. (C) Quantification of CENP-C immunofluorescence from A and B, as indicated. (D-E) Control (top panels) or siLMNA HeLa cells (bottom panels) were immunostaining for CENP-T (green), and ACA (red) with their chromosomes counterstained using DAPI (blue) as indicated. Interphase cells are depicted in D while late telophase cells are depicted in E. (F) Quantification of CENP-T immunofluorescence from D and E, as indicated. (G) Western blot of HeLa cells confirming the generation of *LMNA KO* cell-line using CRISPR technology. Actin is used as a loading control. (H) Freshly thawed *LMNA KO* and WT MEFs were cultured for 15 weeks and the levels of kinetochore CENP-T was quantified at the time points as indicated in the graph. (I) RNA sequencing analysis of WT and *LMNA KO* MEFs depicted as a volcano plot showing the 176 upregulated genes (red) and the 374 downregulated genes (blue). (J) Western blot showing endogenous levels of CENP-A and CENP-C proteins in HeLa cells subjected to BANF1, or EMD knockdown, or to *LMNA KO*. Scale bars, 5 µm.

**Supplementary Figure 3. Depletion of Lamin A-associated proteins (LAAPs), BANF1 and EMD, interferes with centromere assembly.** (A-B) Control interphase (A) or metaphase (B) HeLa cells (top panels) as well as those treated with siBANF1 (middle panels) or siEMD (bottom panels) were immunostained for CENP-A (green) and the chromosomes were counterstained using DAPI (blue). (C) Quantification of CENP-A immunofluorescence after siBANF1 (top) or siEMD (bottom) in the two conditions depicted in A and B. (D-E) Similar to the conditions in A and B but CENP-C (green) was detected in this case instead of CENP-A. (F) Quantification of CENP-C immunofluorescence after siBANF1 (top) or siEMD (bottom) in the two conditions depicted in D and E. (G-H) Similar to the conditions in A and B but CENP-T (green) was detected in this case instead of CENP-A. (I) Quantification of CENP-T immunofluorescence after siBANF1 (top) or siEMD (bottom) in the two conditions depicted in G and H. Scale bars, 5 µm.

**Supplementary Figure 4. Outer kinetochore assembly and mitotic progression is severely perturbed after the inhibition of LAAPs.** (A) Prometaphase HeLa cell controls or those treated with siBANF1 were immunostained for BUB1 (green) with the chromosomes counterstained using DAPI (blue). (B) Prometaphase *WT* or *LMNA KO* MEFs were immunostained for BUB1 (green) with the chromosomes counterstained using DAPI (blue). (C) Quantification of BUB1 immunofluorescence in the conditions as depicted in A and B. (D) Metaphase HeLa cells treated with control or siEMD were immunostained for ZWINT-1 (green) with the chromosomes counterstained using DAPI. (E) Same conditions as in D but in RPE1 cells. (F) Quantification of ZWINT-1 immunofluorescence the conditions as depicted in D and E. (G) Quantification of mitotic progression depicting the percentage of mitotic cells in different stages of mitosis in *LMNA (labelled as LA) KO* HeLa cells. (H) Graph from cells in G depicting the percent of mitotic cells in anaphase that exhibited chromosome mis-segregation. (I-J) Same quantification as described in G and H but with control or siBANF1 cells. (K-L) Same quantification as described in G and H but with control or siEMD cells. Scale bars, 5 µm.

## Supplemental Video Legends

**Video S1. GFP-BANF1 localizes to the chromosomal core domains within minutes after the anaphase onset.**

Live images of HeLa cells stably expressing GFP-tagged BANF1 and the chromosomes labeled with DNA dye, Hoechst, were captured every 30 sec, for a total period of ∼ 30 min. The movie was sped up to ∼ 120 x and played at 4 frames/s. Scale bars, refer to Figure S1C.

**Video S2. WT GFP-LMNA localizes normally to the chromosomal core domains.**

Live images of HeLa cells stably expressing Histone H2B-RFP (to mark the chromosomes) and transfected with a GFP-tagged WT LMNA construct, were captured every 1 min, for a total period of ∼ 25 min. The movie was sped up to ∼ 240 x and played at 4 frames/s. Scale bars, refer to Figure 1F.

**Video S3. E145K GFP-LMNA does not localize properly to the chromosomal core domains.**

Live images of HeLa cells stably expressing Histone H2B-RFP (to mark the chromosomes) and transfected with a GFP-tagged E145K mutant LMNA construct, were captured every 1 min, for a total period of ∼ 30 min. The movie was sped up to ∼ 240 x and played at 4 frames/s. Scale bars, refer to Figure 1F.

## References

1. Takeuchi, K., and Fukagawa, T. (2012). Molecular architecture of vertebrate kinetochores. Experimental cell research 318, 1367–1374.

2. Hori, T., and Fukagawa, T. (2012). Establishment of the vertebrate kinetochores. Chromosome Res 20, 547–561.

3. Zasadzińska, E., and Foltz, D.R. (2017). Orchestrating the Specific Assembly of Centromeric Nucleosomes. Prog Mol Subcell Biol 56, 165–192.

4. Ray-Gallet, D., and Almouzni, G. (2021). The Histone H3 Family and Its Deposition Pathways. Adv Exp Med Biol 1283, 17–42.

5. Cheeseman, I.M., and Desai, A. (2008). Molecular architecture of the kinetochore-microtubule interface. Nat Rev Mol Cell Biol 9, 33–46.

6. Musacchio, A., and Desai, A. (2017). A Molecular View of Kinetochore Assembly and Function. Biology (Basel) 6.

7. Varma, D., and Salmon, E.D. (2012). The KMN protein network--chief conductors of the kinetochore orchestra. J Cell Sci 125, 5927–5936.

8. Güttinger, S., Laurell, E., and Kutay, U. (2009). Orchestrating nuclear envelope disassembly and reassembly during mitosis. Nature Reviews Molecular Cell Biology 10, 178–191.

9. Liu, S., and Pellman, D. (2020). The coordination of nuclear envelope assembly and chromosome segregation in metazoans. Nucleus 11, 35–52.

10. Torras-Llort, M., Medina-Giró, S., Escudero-Ferruz, P., Lipinszki, Z., Moreno-Moreno, O., Karman, Z., Przewloka, M.R., and Azorín, F. (2020). A fraction of barrier-to-autointegration factor (BAF) associates with centromeres and controls mitosis progression. Communications biology 3, 454.

11. Shin, J.Y., and Worman, H.J. (2022). Molecular Pathology of Laminopathies. Annu Rev Pathol 17, 159–180.

12. Taimen, P., Pfleghaar, K., Shimi, T., Möller, D., Ben-Harush, K., Erdos, M.R., Adam, S.A., Herrmann, H., Medalia, O., Collins, F.S., et al. (2009). A progeria mutation reveals functions for lamin A in nuclear assembly, architecture, and chromosome organization. Proceedings of the National Academy of Sciences of the United States of America 106, 20788–20793.

13. Haraguchi, T., Koujin, T., Hayakawa, T., Kaneda, T., Tsutsumi, C., Imamoto, N., Akazawa, C., Sukegawa, J., Yoneda, Y., and Hiraoka, Y. (2000). Live fluorescence imaging reveals early recruitment of emerin, LBR, RanBP2, and Nup153 to reforming functional nuclear envelopes. J Cell Sci 113 (Pt 5), 779–794.

14. Haraguchi, T., Koujin, T., Segura-Totten, M., Lee, K.K., Matsuoka, Y., Yoneda, Y., Wilson, K.L., and Hiraoka, Y. (2001). BAF is required for emerin assembly into the reforming nuclear envelope. J Cell Sci 114, 4575–4585.

15. Haraguchi, T., Kojidani, T., Koujin, T., Shimi, T., Osakada, H., Mori, C., Yamamoto, A., and Hiraoka, Y. (2008). Live cell imaging and electron microscopy reveal dynamic processes of BAF-directed nuclear envelope assembly. J Cell Sci 121, 2540–2554.

16. Samwer, M., Schneider, M.W.G., Hoefler, R., Schmalhorst, P.S., Jude, J.G., Zuber, J., and Gerlich, D.W. (2017). DNA Cross-Bridging Shapes a Single Nucleus from a Set of Mitotic Chromosomes. Cell 170, 956–972.e923.

17. Liu, J., Lee, K.K., Segura-Totten, M., Neufeld, E., Wilson, K.L., and Gruenbaum, Y. (2003). MAN1 and emerin have overlapping function(s) essential for chromosome segregation and cell division in Caenorhabditis elegans. Proceedings of the National Academy of Sciences of the United States of America 100, 4598–4603.

18. Eisch, V., Lu, X., Gabriel, D., and Djabali, K. (2016). Progerin impairs chromosome maintenance by depleting CENP-F from metaphase kinetochores in Hutchinson-Gilford progeria fibroblasts. Oncotarget 7, 24700–24718.

19. Montes de Oca, R., Lee, K.K., and Wilson, K.L. (2005). Binding of barrier to autointegration factor (BAF) to histone H3 and selected linker histones including H1.1. The Journal of biological chemistry 280, 42252–42262.

20. Jansen, L.E., Black, B.E., Foltz, D.R., and Cleveland, D.W. (2007). Propagation of centromeric chromatin requires exit from mitosis. J Cell Biol 176, 795–805.

21. Hayashi, T., Fujita, Y., Iwasaki, O., Adachi, Y., Takahashi, K., and Yanagida, M. (2004). Mis16 and Mis18 are required for CENP-A loading and histone deacetylation at centromeres. Cell 118, 715–729.

22. Fujita, Y., Hayashi, T., Kiyomitsu, T., Toyoda, Y., Kokubu, A., Obuse, C., and Yanagida, M. (2007). Priming of centromere for CENP-A recruitment by human hMis18alpha, hMis18beta, and M18BP1. Dev Cell 12, 17–30.

23. Maddox, P.S., Hyndman, F., Monen, J., Oegema, K., and Desai, A. (2007). Functional genomics identifies a Myb domain-containing protein family required for assembly of CENP-A chromatin. J Cell Biol 176, 757–763.

24. Foltz, D.R., Jansen, L.E., Bailey, A.O., Yates, J.R., 3rd, Bassett, E.A., Wood, S., Black, B.E., and Cleveland, D.W. (2009). Centromere-specific assembly of CENP-a nucleosomes is mediated by HJURP. Cell 137, 472–484.

25. Dunleavy, E.M., Roche, D., Tagami, H., Lacoste, N., Ray-Gallet, D., Nakamura, Y., Daigo, Y., Nakatani, Y., and Almouzni-Pettinotti, G. (2009). HJURP is a cell-cycle-dependent maintenance and deposition factor of CENP-A at centromeres. Cell 137, 485–497.

26. Nardi, I.K., Zasadzińska, E., Stellfox, M.E., Knippler, C.M., and Foltz, D.R. (2016). Licensing of Centromeric Chromatin Assembly through the Mis18α-Mis18β Heterotetramer. Molecular cell 61, 774–787.

27. Pan, D., Klare, K., Petrovic, A., Take, A., Walstein, K., Singh, P., Rondelet, A., Bird, A.W., and Musacchio, A. (2017). CDK-regulated dimerization of M18BP1 on a Mis18 hexamer is necessary for CENP-A loading. Elife 6.

28. Black, B.E., Jansen, L.E.T., Maddox, P.S., Foltz, D.R., Desai, A.B., Shah, J.V., and Cleveland, D.W. (2007). Centromere Identity Maintained by Nucleosomes Assembled with Histone H3 Containing the CENP-A Targeting Domain. Molecular cell 25, 309–322.

29. Moree, B., Meyer, C.B., Fuller, C.J., and Straight, A.F. (2011). CENP-C recruits M18BP1 to centromeres to promote CENP-A chromatin assembly. J Cell Biol 194, 855–871.

30. Stellfox, M.E., Nardi, I.K., Knippler, C.M., and Foltz, D.R. (2016). Differential Binding Partners of the Mis18α/β YIPPEE Domains Regulate Mis18 Complex Recruitment to Centromeres. Cell Rep 15, 2127–2135.

31. Kim, Y., and Zheng, Y. (2013). Generation and characterization of a conditional deletion allele for Lmna in mice. Biochem Biophys Res Commun 440, 8–13.

32. McKinley, K.L., and Cheeseman, I.M. (2017). Large-Scale Analysis of CRISPR/Cas9 Cell-Cycle Knockouts Reveals the Diversity of p53-Dependent Responses to Cell-Cycle Defects. Dev Cell 40, 405–420.e402.

33. Chang, W., Folker, E.S., Worman, H.J., and Gundersen, G.G. (2013). Emerin organizes actin flow for nuclear movement and centrosome orientation in migrating fibroblasts. Mol Biol Cell 24, 3869–3880.

34. Ostlund, C., Ellenberg, J., Hallberg, E., Lippincott-Schwartz, J., and Worman, H.J. (1999). Intracellular trafficking of emerin, the Emery-Dreifuss muscular dystrophy protein. J Cell Sci 112 (Pt 11), 1709–1719.

35. Lee, M.S., and Craigie, R. (1998). A previously unidentified host protein protects retroviral DNA from autointegration. Proceedings of the National Academy of Sciences of the United States of America 95, 1528–1533.

36. Rahi, A., Chakraborty, M., Agarwal, S., Vosberg, K.M., Agarwal, S., Wang, A.Y., McKenney, R.J., and Varma, D. (2023). The Ndc80-Cdt1-Ska1 complex is a central processive kinetochore-microtubule coupling unit. J Cell Biol 222.

37. Walstein, K., Petrovic, A., Pan, D., Hagemeier, B., Vogt, D., Vetter, I.R., and Musacchio, A. (2021). Assembly principles and stoichiometry of a complete human kinetochore module. Sci Adv 7.

