## Supplementary figures and images for "Nuclear lamin A-associated proteins are required for centromere assembly"

### Supplemental Figures

Supplemental Figure 1

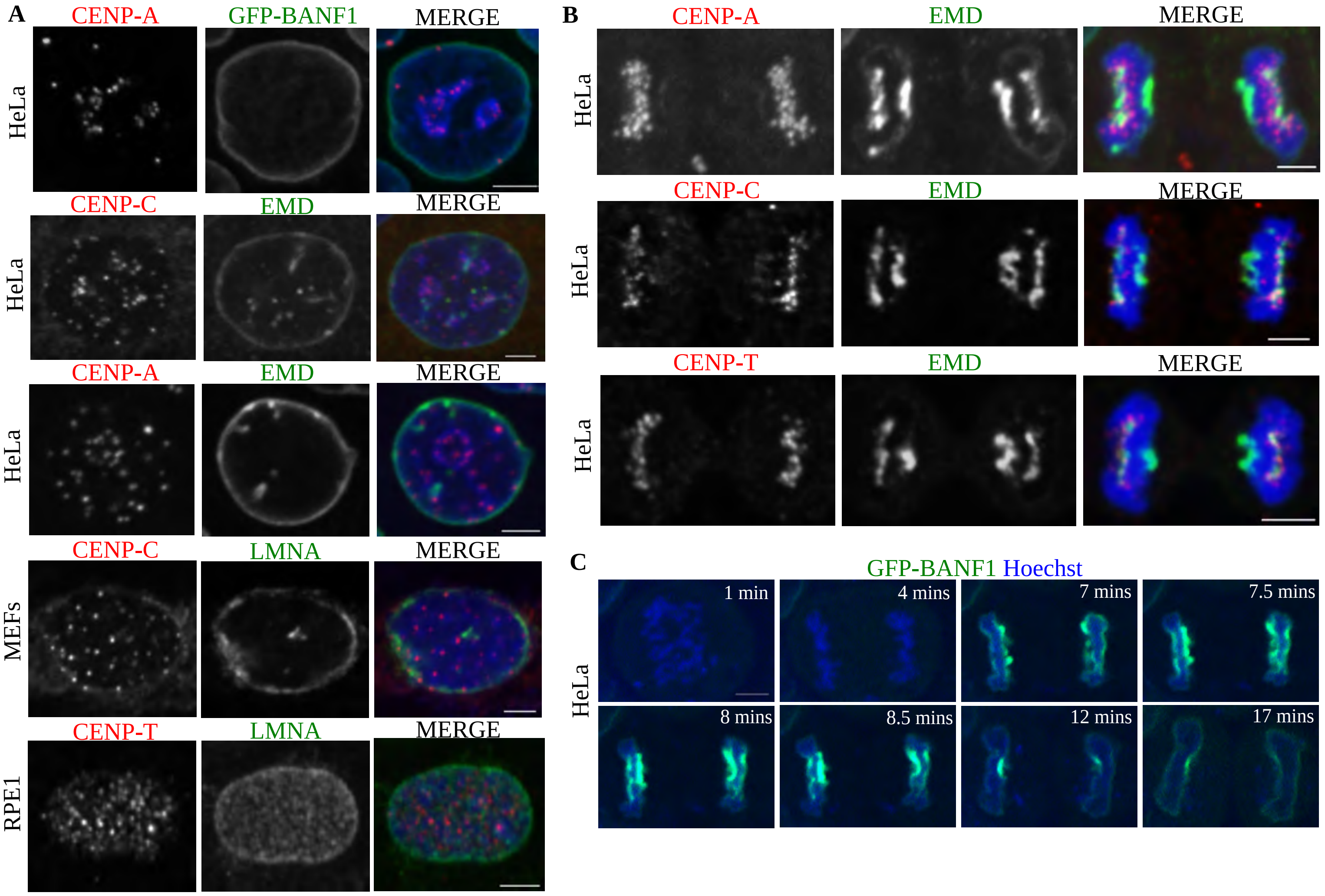

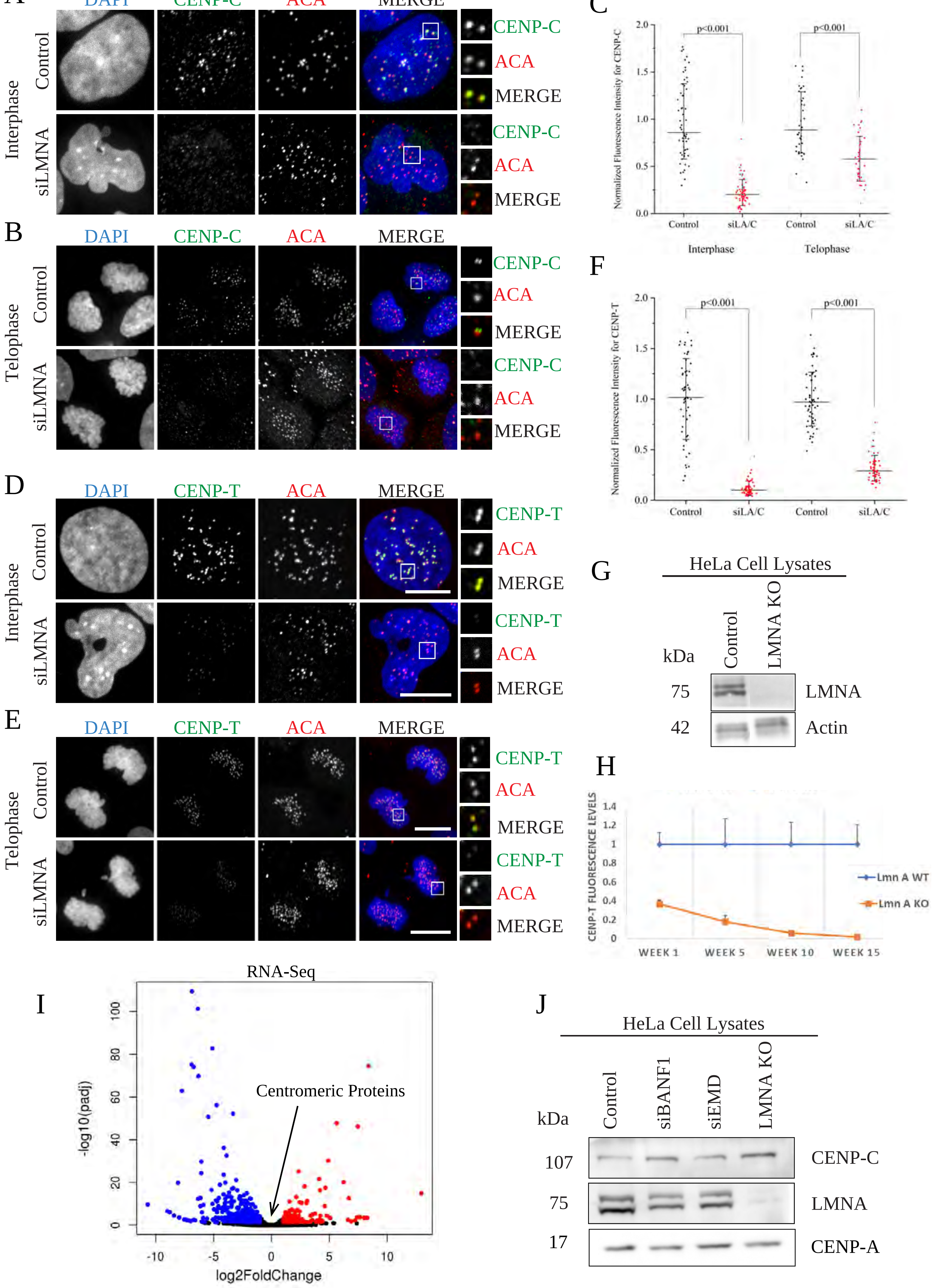

#### Supplemental Figure 3

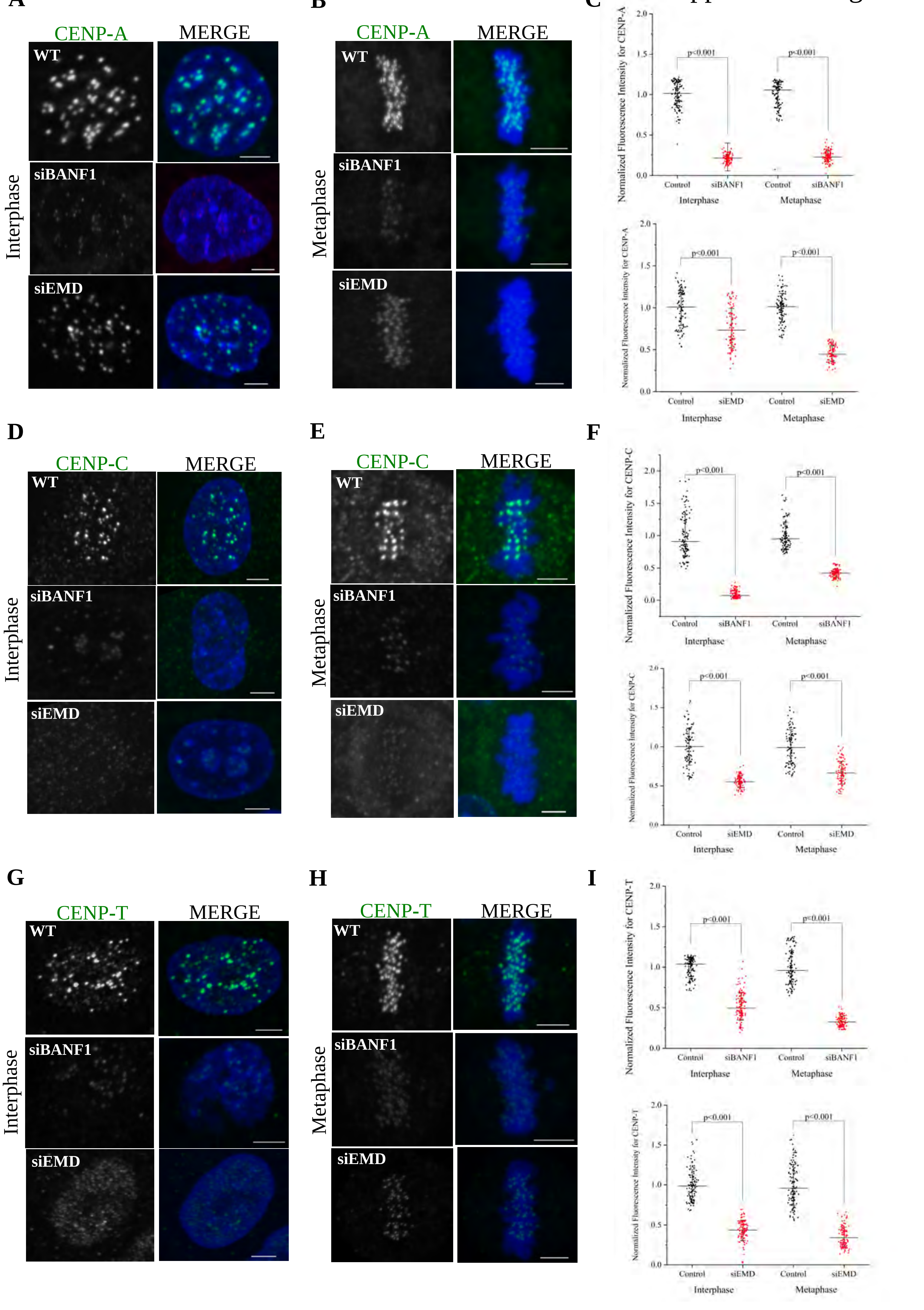

Supplemental Figure 4

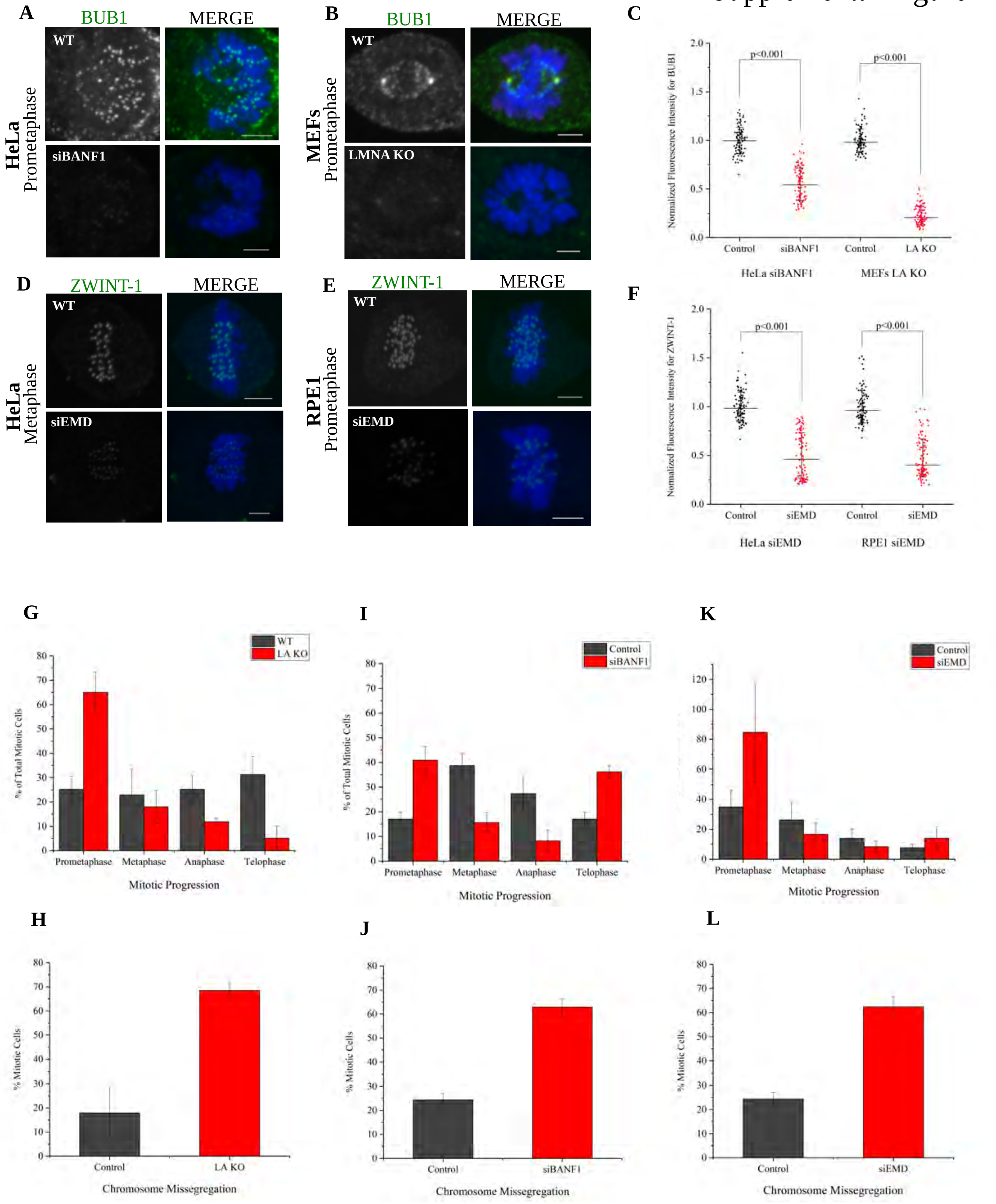
